# Circuits in the absence of cortical layers: increased callosal connectivity in reeler mice revealed by brain-wide input mapping of VIP neurons in barrel cortex

**DOI:** 10.1101/2020.04.19.048868

**Authors:** Georg Hafner, Julien Guy, Mirko Witte, Pavel Truschow, Alina Rüppel, Nikoloz Sirmpilatze, Rakshit Dadarwal, Susann Boretius, Jochen F. Staiger

## Abstract

The neocortex is composed of layers. Whether layers constitute an essential framework for the formation of functional circuits is not well understood. We investigated if neurons require the layer organization to be embedded into brain-wide circuits using the reeler mouse. This mutant is characterized by a migration deficit of cortical neurons so that no layers are formed. Still, neurons retain their properties and reeler mice show little cognitive impairment. We focused on VIP neurons because they are known to receive strong long-range inputs and have a typical laminar bias towards upper layers. In reeler these neurons are more distributed across the cortex. We mapped the brain-wide inputs of VIP neurons in barrel cortex of wildtype and reeler mice with rabies virus tracing. Innervation by subcortical inputs was not altered in reeler, in contrast to the cortical circuitry. Numbers of long-range ipsilateral cortical inputs were reduced in reeler, while contralateral inputs were strongly increased. Reeler mice had more callosal projection neurons. Hence, the corpus callosum was larger in reeler as shown by structural imaging. We argue that in the absence of cortical layers, circuits with subcortical structures are maintained but cortical neurons establish a different network capable to preserve cognitive functions.

## INTRODUCTION

The neurons of the mammalian neocortex are stacked in layers (L). This organization is considered as an essential scaffolding principle that evolution could expand to generate increasingly sophisticated cognitive abilities (Shepherd and Rowe, 2017). Layers are seen as units of processing and their input-output connectivity defines the flow of information within the cortex (Feldmeyer, 2012; Miller, 2001). However, the notion of layers structuring the flow of cortical processing might be misleading. Cortical computations require foremost a functional neuronal circuit between defined cell types (Guy and Staiger, 2017; Harris and Shepherd, 2015). Because neurons exhibit surprising specificity in targeting of other neurons, there seem to be strict rules of connectivity (Kasthuri et al., 2015; Motta et al., 2019). These wiring rules might run in the framework of layers but perhaps are not determined by it. Whether the organization of layers constitutes an indispensable feature for the formation of cortical circuits is still unclear.

Therefore, we asked the question if correct laminar position is a necessary prerequisite for a neuron to receive inputs and become properly embedded in brain-wide circuits. An excellent model system to study the significance of layers is the reeler mouse (Guy and Staiger, 2017). Homozygous mice lack the protein reelin, which is secreted by Cajal-Retzius cells during development, to orchestrate the migration of neurons during development (Lee and D’Arcangelo, 2016). In consequence, the formation of layers is strongly compromised. In the somatosensory cortex, neurons destined to become part of a certain layer are almost uniformly dispersed across the cortex (Boyle et al., 2011; Dekimoto et al., 2010; Wagener et al., 2010). However, this perturbation seems to have little effect on properties of neurons and formation of functional circuits. Neurons retain their molecular fate as well as their electrophysiological properties (Guy et al., 2016; Silva et al., 1991; Wagener et al., 2010). The structure of synaptic boutons remains virtually unchanged (Prume et al., 2018, 2019). Areal boundaries within the cortex are preserved (Boyle et al., 2011). Moreover, whisker stimulation of reeler mice leads to the same hemodynamic responses in the cortex and activates the same cell types indicating a preservation of circuit modules comparable if not identical to a layered cortex (Guy et al., 2015; Wagener et al., 2016). For these reasons, the reeler mouse seems ideally suited to study functional brain circuitries of defined cell types in the absence of layers.

We decided to focus on the connectivity of vasoactive intestinal polypeptide (VIP) expressing neurons. They constitute a major subgroup of GABAergic inhibitory neurons (Tremblay et al., 2016). There are several reasons why we chose this type of neuron. First, VIP neurons have a very distinct laminar distribution with about 60% occurring in LII/III, 20% in LIV and only very few cells in deep layers (Prönneke et al., 2015). So they are heavily biased towards upper layers. Second, VIP neurons have been suggested as major integrators of long-range input. They receive more inputs per cell than other inhibitory cell types (Wall et al., 2016). They are most strongly activated by cortical long-range input compared to pyramidal, somatostatin- and parvalbumin expressing neurons (Lee et al., 2013; Zhang et al., 2014). Because VIP cells strongly inhibit somatostatin cells (Walker et al., 2016), which themselves inhibit pyramidal cells (Zhou et al., 2020), it has been proposed that VIP neurons become activated by long-range input during active states, disinhibit pyramidal cells and thereby open a precisely timed window for integration and plasticity at excitatory synapses (Fu et al., 2014; Pfeffer et al., 2013; Williams and Holtmaat, 2019). Third, the input connectivity of VIP neurons is already well characterized across many cortical areas (Ährlund-Richter et al., 2019; Sun et al., 2019; Wall et al., 2016; Zhang et al., 2016). Finally, with the VIP-Cre line we can access these cells with high specificity having no contamination from other cell types (Prönneke et al., 2015; Taniguchi et al., 2011).

We first confirmed that VIP neurons lose their laminar bias in reeler mice, establishing this mouse as a valid model system to study the circuitry of VIP cells in the absence of layers. Then we generated maps of brain-wide long-range input to VIP neurons in the barrel cortex of wildtype (WT) and reeler mice, using rabies virus tracing. While we found that subcortical inputs innervated malpositioned VIP cells to the same extent as in a layered cortex, the balance of cortical inputs was fundamentally different. Ipsilateral cortical input was reduced, contralateral cortical input was increased compared to WT. Reeler mice had more callosal projecting neurons (CPNs) and hence a larger corpus callosum. We argue that the absence of layers has little effect on innervation by subcortical structures but induces a different circuit arrangement within the cortex.

## MATERIAL AND METHODS

### Experimental animals

We crossed the reeler line (B6C3Fe a/a-Relnrl/J, The Jackson Laboratory, Bar Harbor, USA) with the VIP-Cre line (VIPtm1(cre)Zjh, The Jackson Laboratory) to breed VIP-Cre/reeler mice heterozygous for reelin mutation and homozygous for Cre. These animals were crossed to generate VIP-Cre/reeler mice homozygous for reelin knockout. WT littermates or animals from the VIP-Cre line were used for comparison in tracing experiments. For control experiments to check the quality of our Cre-dependent constructs, we used C57BL/6J wildtype mice (The Jackson Laboratory). For tracing experiments not requiring Cre expression and for imaging we used WT and homozygous littermate pairs of the reeler line.

To visualize the population of VIP cells, we crossed the VIP-Cre/reeler line with the Ai9 tdTomato reporter line (B6.Cg-Gt(ROSA)26Sortm9(CAG-tdTomato)Hze/J, The Jackson Laboratory) to achieve tdTomato expression in VIP neurons.

Mice were housed in standard cages in a 12h light/dark cycle and with ad libitum access to food and water. All experimental procedures were performed in accordance with German laws on animal research (TierSchG und TierSchVersV 2013). All tracing experiments were performed with 12-20 weeks-old mice of either gender.

### Viral constructs

pAAV-DIO-TVA^66T^-EGFP-oG was generated based on the backbone of pAAV-hSyn-oG-EGFP-TVA-WPRE-hGpA (gift from Euiseok Kim). The sequence for oG-EGFP was extracted using PCR with the primers F1-CTATACGAAGTTATGGTACCTTAGAGCCG; R1-GGATCCGGAGCTACTAACTTCAGC. The sequence for TVA^66T^ was extracted from pAAV-CAG-FLEX-TC^66T^ ((Miyamichi et al., 2013), Addgene #48331) using PCR with primers F1-AGTAGCTCCGGATCCCCCACCCCCCTTGGATGC; R1-CGGTAACGTGACCGGTAACGG. pAAV backbone was cut with restriction enzymes KpnI-HF and AgeI-HF to open the backbone and oG-EGFP fragment was reinserted together with the TVA^66T^ fragment using infusion cloning with an In-Fusion HD cloning kit (Takara bio). The plasmid was packaged into AAV8 by the SALK Viral Vector Core, from where it is available.

For RV-tracing experiments, 100-200 nl of AAV8-DIO-TVA^66T^-EGFP-oG was injected at a titer of 1.6*10^13 IU/ml. RV-SADΔG-mCherry (EnvA) was injected at the same location fourteen days later at a titer of 1*10^7 IU/ml. Animals were sacrificed seven days later. For non-transsynaptic tracing experiments 150 nl of AAV-retrograde-hSyn-EGFP (Addgene, #50465) at a titer of 2-4*10^12 IU/ml was used. Animals were sacrificed after fourteen days.

### Surgery and viral injection

For viral tracing experiments, mice underwent intrinsic signal optical imaging to localize the whisker C2 related cortical column. The injection pipette was inserted at this location in WT and reeler animals. The surgery was performed as in (Hafner et al., 2019).

For sedation and analgesia, mice were injected intraperitoneally with 10 μg/g xylazine (Xylariem, Ecuphar) and 0.065 μg/g buprenorphine (Temgesic, Individor UK Limited) in sterile saline, respectively. Anesthesia was induced with 3% isoflurane (vol/vol) and maintained between 0.5 and 1% throughout the entire surgical procedure (Harvard Apparatus, USA). Mice were mounted on a custom-built frame with rigid earbars. A mixture of 2μg/g bupivacaine/lidocaine (Astra Zeneca) was injected subcutaneously under the scalp for local anesthesia. A heating pad was used to maintain body temperature at 37 °C (ATC 1000, World Precision Instruments, Florida). Subsequently, a small incision was made in the scalp to expose the right hemisphere of the skull. The bone over the somatosensory area was thinned to transparency with a dental drill (OS-40, Osada Electric Company, Japan). Then, the location of the C2 whisker-related column was determined and mapped on the blood vessel pattern as described below. The bone above the target area was removed with a syringe tip. A glass pipette cut to 20 μm tip diameter (Drummond Scientific Co, USA) was front-filled with AAV helper virus. The pipette holder was attached to a micromanipulator (Luigs & Neumann, Germany). The pipette was inserted at the target location into the brain in an approximately 45° angle, orthogonal to the curvature of the cortex. AAV was pressure-injected with a syringe at three depths (750 μm, 500 μm, and 250 μm below pia). To reduce backflow, the needle was left in place at each depth for at least three minutes. The scalp was sutured and the mouse received a subcutaneous injection of 5 μg/g Carprofen (Pfizer) for prolonged pain relief. Fourteen days later, the mouse was injected with RV-mCherry without prior imaging. The injection was guided based on the blood vessel pattern and landmarks from the previous surgery.”

### Intrinsic signal optical imaging

This procedure was performed as described earlier (Guy et al., 2015; Hafner et al., 2019). Briefly, Whisker C2 on the left muzzle was stimulated with a piezo actuator at 5Hz. Red light was shone on the cortex exposed, thinned cortex above the right barrel field and its reflectance was measured with a CCD camera. Changes in reflectance were average across 30 trials. The blood vessel pattern and the intrinsic signal were overlaid to guide the subsequent injection.

### Fixation and tissue sectioning

Mice were sacrificed by injecting an overdose of ketamine (100 μg/g; Medistar) and perfused transcardially with ice-cold 10% sucrose solution followed by 4% paraformaldehyde (PFA) in 0.1M phosphate buffer. The brain was postfixed in 4% PFA for 4h. A vibratome (VT 1200 S, Leica) was used to section the brain into 100 μm-thick coronal sections rostral and caudal to barrel cortex, while barrel cortex was sectioned at 50 μm-thick intervals. Sections spanning the barrel cortex were subjected to immunohistochemistry while all other sections were stained for 4’,6-diamidino-2-phenylindole (DAPI) only.

### Immunohistochemistry

Barrel cortex sections were washed in TRIS buffer (TB) 1×15 min, TRIS-buffered saline (TBS) 1×15 min and TBS + 0.5% Triton X-100 (TBST) 2×15 min, all at pH 7.6. For blocking, sections were incubated 90 min at room temperature in 0.25% bovine serum albumin/10% goat serum/TBST (Jackson Immuno Research). For primary antibody labeling, sections were incubated 48-72 h at 4°C with (i) chicken anti-GFP (Aves) diluted 1:500, (ii) mouse anti-RFP (Rockland) diluted 1:2000. Sections were washed 4×15 min with TBST. For secondary antibody labeling, sections were incubated 4h at room temperature with (i) Alexa Fluor 488-conjugated goat anti-chicken IgG (Molecular Probes), (ii) Alexa Fluor 568-conjugated goat anti-mouse IgG2a (Molecular Probes) diluted 1:500 in TBST. In some cases, sections were stained in addition with primary antibody guinea pig anti-vGluT2 (Millipore) diluted 1:2000 and secondary antibody Alexa Fluor 633-conjugated goat anti-guinea pig (Molecular probes) diluted 1:500. After washing 2×15 min with TBST and 1×15 min with TBS, sections were stained with DAPI, diluted 1:1000 in TBS. After several washes in TB, sections were mounted in Aqua-Poly-Mount (Polysciences).

### Fluorescent *in-situ* hybridization

Sections of the VIPCre/Reeler/tdTomato mouse were stained with a riboprobe against VIP RNA. VIP riboprobe was generated as described earlier (Prönneke et al., 2015) using the following primers FP: CCTGGCA TTCCTGATACTCTTC; RP: ATTCTCTGATTTCAGCTCTGCC (527 bp; Allen Brain Atlas Riboprobe ID: RP_070116_02_E09).The staining procedure was performed exactly as in (Prönneke et al., 2015). The tdTomato signal was amplified after the in-situ hybridization using the immunohistochemistry protocol described above but without Triton X-100. Primary antibody rabbit anti-RFP (Rockland) was combined with secondary antibody Alexa Fluor 594-conjugated goat anti-rabbit (Molecular probes).

### Image acquisition and processing

Images were acquired on an inverted epifluorescence slide-scanning microscope (Axio Observer, Zeiss) with a 10x objective (NA=0.3) or on 25x objective (NA=0.8). For overview images only one plane was imaged, for the injection sites stacks were acquired. Tiles were stitched after imaging and stacks were deconvoluted in ZEN Blue software (Zeiss) to reduce out-of-focus light. Confocal acquisitions were taken on a LSM 880 (Zeiss) with a 25x objective operated by ZEN black software (Zeiss).

### Magnetic resonance imaging (MRI) data acquisition and preprocessing

The *ex vivo* brain samples (n=3 per group) were imaged in a high-field 9.4 Tesla MR system, using a mouse brain 4-channel coil array (Bruker BioSpin MRI GmbH, Ettlingen, Germany). The scanning protocol included Magnetization Transfer (MT) weighted images and diffusion-weighted images. For MT, a 3D Fast Low Angle Shot (FLASH) sequence was used to acquire three datasets: MT-weighted, Proton-Density-weighted, and T_1_-weighted (repetition time 15 ms, echo time 3.2 ms, flip angles [5°, 5°, 25°], 10 averages, voxel size 125 x 125 x 125 μm^3^). These datasets were used to estimate MT saturation (MTsat) according to the method described by Helms *et al* (Helms et al., 2008). Diffusion-weighted images were acquired using a Stejskal-Tanner pulsed gradient spin-echo sequence (repetition time 2000 ms, echo time 23.2 ms, 20 slices, 3 averages, voxel size 125 x 125 x 500 μm^3^, b values 3000 and 6000 s/mm^2^, 30 directions each, 5 b0 images). The diffusion data were preprocessed through denoising (Veraart et al., 2016), correction for eddy current distortions (Andersson and Sotiropoulos, 2016), motion correction (Avants et al., 2011), and bias field correction (Tustison et al., 2010). A Diffusion Tensor model (Basser et al., 1994) was fitted to the preprocessed data and fractional anisotropy (FA) was derived (Garyfallidis et al., 2014). An RGB color-coded FA map was also computed. To ensure that all images were centered at a common origin and oriented in the same way, we registered the individual mouse brains to the digital template of the Allen Mouse Brain Common Coordinate Framework version 3 (Lein et al., 2007). The registration employed six degrees of freedom (three translations and three rotations) to maximize the mutual information between each individual brain and the template (Avants et al., 2011). Since this constitutes a rigid-body transform, the original volumes of the structures were preserved.

### Corpus callosum (CC) and isocortex segmentation

To measure the cross-sectional area of the CC at its midline crossing in a consistent way across animals, we followed a multi-step process involving the computed maps for MTsat and RGB FA. The MTsat maps, which exhibit good grey/white matter contrast, were used to generate white matter masks through simple thresholding (voxels with MTsat > 0.006 were classified as white matter). We subsequently restricted the white matter masks to the five middle sagittal slices (left-to-right thickness 0.625 mm) and further cropped them along the rostrocaudal (5 mm) and the dorsoventral (3.125 mm) directions—thus confining them within a thin midline slab that contained the entire CC crossing. This slab also included several other white matter tracts, some of which (e.g. dorsal fornix) run adjacent to the CC and are challenging to separate based on MTsat alone. We addressed this issue by incorporating information about the orientation of white matter tracts. This was available in the form of the RGB FA map, which is a color-coded representation of the principal diffusion direction at each voxel (Pajevic and Pierpaoli, 1999). Using ITK-SNAP (Yushkevich et al., 2006), we overlaid the RGB FA map on the MTsat image, and segmented the CC manually, excluding voxels with principal diffusion directions other than left-to-right. In reeler mutants, this step proved especially useful for excluding a midline tract with a rostrocaudal orientation, running dorsal to the CC. Using the final CC mask, we computed its midline cross-sectional area by multiplying the number of CC voxels on the midsagittal slice with the 2D area of each voxel (0.125 x 0.125 mm^2^).

The isocortical volume was derived from MTsat images, with the help of the Allen Brain Atlas. First we pooled all isocortical regions of the atlas into a single mask and overlaid it on each individual MTsat image using ITK-SNAP. Then we manually edited the mask for each brain, until it covered the entire cortical thickness, without protruding into extra-cortical areas. The final isocortical volume was computed by multiplying the number of voxels in the edited mask with the 3D volume of each voxel (0.125 x 0.125 x 0.125 mm^3^).

### Quantification and statistical analysis

Mapping of RV-labeled input cells was done by overlaying the tissue section with the corresponding section of the Allen Brain Atlas (http://mouse.brain-map.org/experiment/thumbnails/100048576?image_type=atlasand). Labeled cells on all sections spanning from Bregma +3 to −4.5 mm were counted manually in Neurolucida (MBF Bioscience). Cell counts in an area were either normalized by the total number of starter cells (input magnitude) or by the total number of inputs cells (input fraction).

For quantification of contralateral input labeled with AAV-retro-EGFP, we manually counted all EGFP-positive cells in contralateral barrel cortex on sections from Bregma −0.8 to −2.1 mm as well as all cells in a volume from pia to white matter with a 200 μm diameter around the injection site. Contralateral neurons were then divided by the number of cells at the injection site as normalization to control for different labeling intensities.

For quantification of VIP neurons in VIP-Cre/tdTomato mice, neurons were manually counted on six sections per mouse on a patch of barrel cortex that spanned from pia to white matter and was 1000 μm wide. In the occasional case a section was lost during the tissue preparation, we interpolated for this animal the number of cells to the same volume.

For a layer-independent analysis of neuronal distribution in the cortex, we divided the space between the pia and the white matter boarder in 20 bins of equal size. We counted the number of labeled cells in each of these bins and normalized them by the total count of cells in all bins. Cell counts were exported with Neurolucida Explorer to Excel. R software (www.R-project.org) was used to sort data and perform statistical test with custom-written code. For pairwise comparisons between WT and reeler mice, data were first tested for normality (Shapiro–Wilk test) and equal variance (Barlett test). For normally distributed data with equal variances, we used the Student’s t-test. For non-normally distributed data, we used the Wilcoxen rank-sum test. For normally distributed data with unequal variances, we used the Welch’s t-test. For multiple comparisons, we used the alpha-error adjustment by Holm. All values are given as mean ± SD. Graphs were produced using Origin software (Origin Lab, USA). Adobe Illustrator and InDesign CS6 were used for arrangement of pictures.

## RESULTS

### VIP neurons are uniformly distributed across the cortex in reeler mice

It is known from previous studies in reeler mice that parvalbumin- (Boyle et al., 2011) and somatostatin-positive (Yabut et al., 2007) neurons lose their typical laminar distribution but VIP neurons have not been investigated yet. We generated VIP-Cre/tdTomato/reeler mice to visualize VIP neurons (Figure 1A, A’). While in WT they showed their typical bias towards the upper layers, in reeler mice VIP neurons appeared homogenously distributed throughout the thickness of the cortex (Figure 1B, B’, Figure S1A). The number of VIP cells in barrel cortex stayed approximately the same (Figure S1B). To check if Cre-expressing tdTomato cells are VIP expressing, we stained sections of reeler barrel cortex against VIP-RNA with in-situ hybridization. We found that 99.4±0.8% of tdTomato cells expressed VIP (n=3 mice, 3 sections each; Figure S1C). This is the same level of specificity demonstrated for the VIP-Cre/tdTomato/WT mouse (Prönneke et al., 2015). In sum, VIP neurons in the reeler mouse retain their VIP expression, appear in equal numbers as in WT but show no laminar bias. Hereafter we used the VIP-Cre/reeler mouse to specifically target VIP neurons and investigate if their altered distribution affects their ability to integrate long-range inputs.

**Figure 1:**
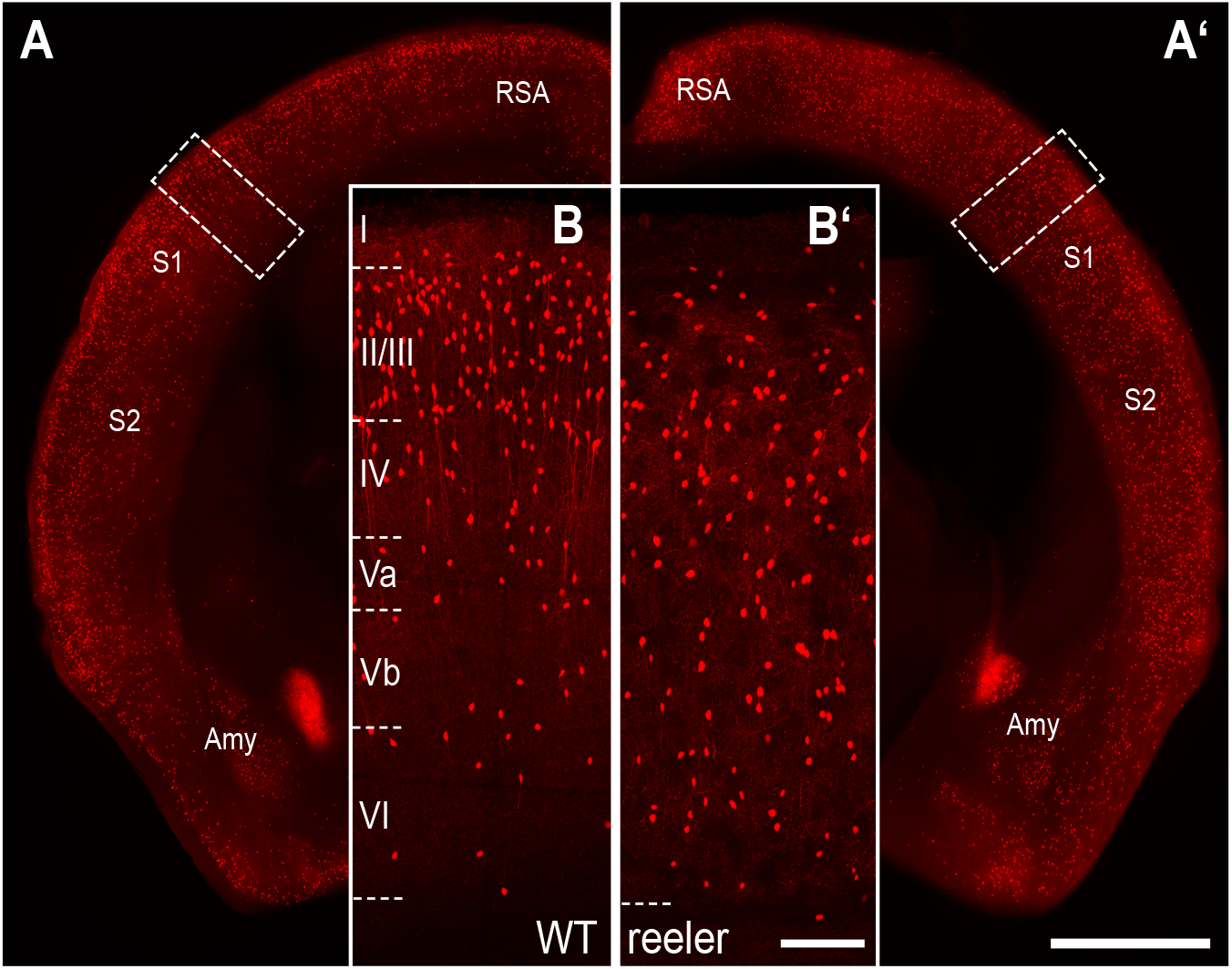
Distribution of VIP cells is different between WT and reeler mice. (A, A’) Coronal section at the level of the barrel cortex of a WT and reeler mouse in which VIP neurons are labeled with tdTomato. The areal/nuclear locations of VIP expression remained the same in WT and reeler (scale bar: 1000 μm; Abbr.: Amy, Amygdala; RSA, retrosplenial agranular cortex; S1/S2, primary/secondary somatosensory cortex). (B, B’) Close-up of cortical tissue in WT and reeler mouse (insert in A, A’). In WT, VIP neurons showed a stronger bias towards upper layers (II-IV). In reeler, VIP neurons were uniformly dispersed across the cortical thickness (scale bar: 100 μm).

### Rabies virus tracing for mapping of brain-wide inputs to VIP neurons

We used Cre-dependent rabies virus tracing, in order to map the monosynaptic long-range inputs to VIP cells in barrel cortex of WT and reeler mice (Wickersham *et al*., 2007; Wall *et al*., 2016; Figure 2). We injected AAV8-DIO-TVA^66T^-EGFP-oG (hereafter AAV-TVA^66T^-EGFP-oG) into the cortex of VIP-Cre mice having WT or reeler genotypes. This lead to the expression of three proteins: the cell surface receptor TVA (here we used a mutated version, TVA^66T^ (Miyamichi et al., 2013)), the optimized version of the rabies glycoprotein (oG; Kim *et al*., 2016) and EGFP (Figure 2A). Modified rabies virus RV-SADΔG-mCherry (EnvA) (hereafter RV-mCherry; Figure 2A) was injected two weeks later at the same location. Because it is coated with the EnvA-ligand TVA, it can only transduce cells presenting TVA on their surface. Furthermore, RV-mCherry is G-deleted, so that it needs the glycoprotein provided in *trans* from the AAV to spread to presynaptic neurons. Its mCherry expression labels these presynaptic neurons red. The starter cells appear yellow due to the mixture of EGFP and mCherry (Figure 2B). This two-virus system allows the visualization of brain-wide monosynaptic inputs to unequivocally identifiable starter cells.

**Figure 2:**
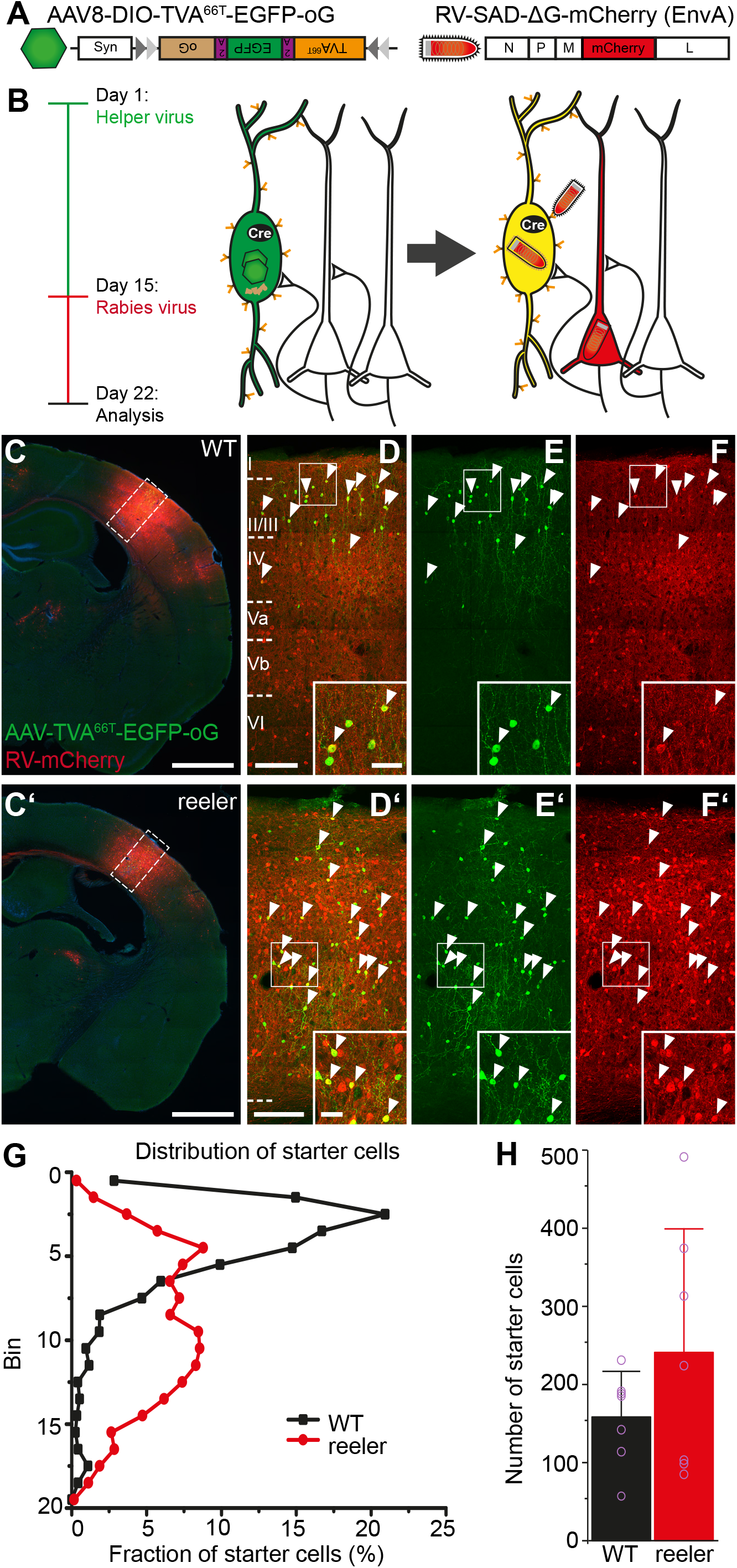
RV-tracing in VIP-Cre mice is based on a different distribution and number of VIP starter cells in WT and reeler mice. (A) Viral constructs for RV-tracing. TVA^66T^ is a mutated version of TVA, to which EnvA in the RV envelope has a reduced affinity. This ensures that low level Cre-independent TVA expression does not permit RV entry into cells. oG is the optimized rabies glycoprotein necessary for transsynaptic spread. (B) Injection of Cre-dependent helper AAV on day 1 induces high expression of TVA^66T^ and oG only in VIP cells. RV-mCherry pseudotyped with TVA-ligand EnvA is injected 14 days later to infect VIP neurons and spread from there to first-order presynaptic neurons using oG. Seven days later, starter VIP cells appear yellow due to the mixture of fluorophores, presynaptic neurons have solely mCherry. (C, C’) Coronal sections through an injection site in the barrel cortex of WT and reeler mice (scale bar: 1000 μm). (D/D’-F/F’) Inserts in C, C’. Cells marked by white arrowheads are double-labeled starter cells that have been co-transduced by AAV-TVA^66T^-EGFP-oG and RV-mCherry. Inserts at the bottom show some of these cells in higher resolution. Exclusively RV-mCherry-positive cells represent local inputs, presynaptic to the starter cells, forming a dense network of cell bodies and neuropil (scale bar overview: 100 μm; scale bar insert: 20 μm). (G) Distribution of starter cells across the cortical depth. Cortical thickness was divided into 20 equal-sized bins. The proportion of starter cells in each bin was plotted. While in WT starter cells were predominantly in the upper third, starter cells in reeler were much more dispersed. (H) Number of starter cells in each genotype. Purple circles represent individual animals. (n=7 per group; mean ± SD).

To check if our viruses are specific, we performed control experiments in BL6 animals without Cre. Injection of RV-mCherry alone did not result in any labeling, showing that TVA is required for RV entry into cells (Figure S1A). When we injected AAV-TVA^66T^-EGFP-oG and RV-mCherry as in tracing experiments, we did not detect any RV-mCherry labeling, neither at the injection site nor in the thalamus (Figure S1B). A few EGFP-positive cells most likely resulted from a minimal Cre-independent expression of the AAV construct. While RV requires very little TVA to enter cells, which can result in Cre-independent leak expression, TVA^66T^ has lower affinity to RV and thus needs to be expressed in much higher levels (Miyamichi et al., 2013). Therefore, this receptor abolishes leak expression and each mCherry-positive neuron can be surely considered an input neuron.

### Injections were centered on the whisker C2 representation

In the barrel cortex, each whisker is represented by a cortical column. Although the reeler barrel cortex has no laminar organization, it retains a somatotopic organization such that adjacent whiskers are represented by adjacent modules (Guy et al., 2015). We localized the cortical whisker C2 representation to guide our injections using intrinsic signal optical imaging (Grinvald et al., 1986; Guy et al., 2015). Repetitive single-whisker C2 stimulation elicited hemodynamic responses, which appeared similar among WT and reeler mice (Figure S2A, C). The overlay of the signal with the blood vessel pattern on the cortical surface (Figure S2B, D) provided a map to accurately target the whisker-related cortical module with an injection (Hafner et al., 2019). As a proof of principle, we injected AAV-TVA^66T^-EGFP-oG and RV-mCherry centered on C2 in a WT animal and sectioned the cortex tangentially. We could confirm our target specificity as the highest density of inputs was present within C2 (Figure S2E). In this way we could target VIP neurons that belong to the same functional modules in WT and reeler mice to have optimal comparability between genotypes.

### VIP starter cells in reeler did not show a laminar bias

VIP-Cre and VIP-Cre/reeler mice (n = 7 each group) were injected with AAV-TVA^66T^-EGFP-oG and two weeks later with RV-mCherry into the right barrel cortex. Cells double-labeled with EGFP/mCherry were considered starter cells and counted on each section where they appeared (Figure 2C-F). To visualize the distribution of starter cells in the two genotypes, we divided the cortex in 20 equal-sized bins and calculated the proportion of starter cells in each bin relative to the total number of starter cells (Figure 2G). The starter cell distribution matched the general distribution of VIP cells in both genotypes. While in WT there was a clear bias of starter cells towards the upper third of the cortex, the distribution of starter cells in reeler was broader with the majority of cells in the middle part of the cortex. On average there was a higher number of starter cells in reeler (Figure 2H; mean WT: 158.3±58.3; mean reeler: 240.9±158.1; Wilcoxon rank sum test, W=30, p= 0.54). This difference could arise because brain tissue in reeler might absorb virus solution better. The potential confounder of such a bias is addressed below.

### VIP cells received input from the same areas in WT and reeler

To address the question if the absence of layers impacts the capacity of VIP cells to receive proper long-range inputs, we manually counted all RV-labeled cells in the entire brain but omitted cells in the barrel cortex itself because of their extreme abundance. Therefore, local connectivity of VIP neurons was not assessed. Each coronal section was overlaid with the corresponding atlas section of the Allen Brain atlas. RV-mCherry-positive cells were assigned to an area, based on the outlines of the atlas and the cytoarchitectonic features discernable with nuclear stain.

VIP neurons in reeler mice received input from the very same areas as WT mice with no exception. Examples for areas in which we found transynaptically labeled cells are presented in Figure 3. Secondary somatosensory cortex contributed the highest number of input cells in WT and reeler (Figure 3B, B’). In some areas morphological differences of input cells became salient, for example in motor cortex where somata appeared larger (Figure 3A, A’). Quantitative differences were easily noticeable in contralateral barrel cortex, which contained more cells in reeler (Figure 3C, C’) and primary auditory cortex, which contained fewer cells in reeler compared to WT (Figure 3D, D’). Thalamic inputs appeared very similar (Figure 3E, E’). In visual cortex the distribution of projection neurons was clearly different with inputs in WT located at the LIII/IV border but being more dispersed in reeler (Figure 3F, F’). From these observations we suspected that the absence of layers could affect most strongly the magnitude and the proportion of inputs to VIP cells.

**Figure 3:**
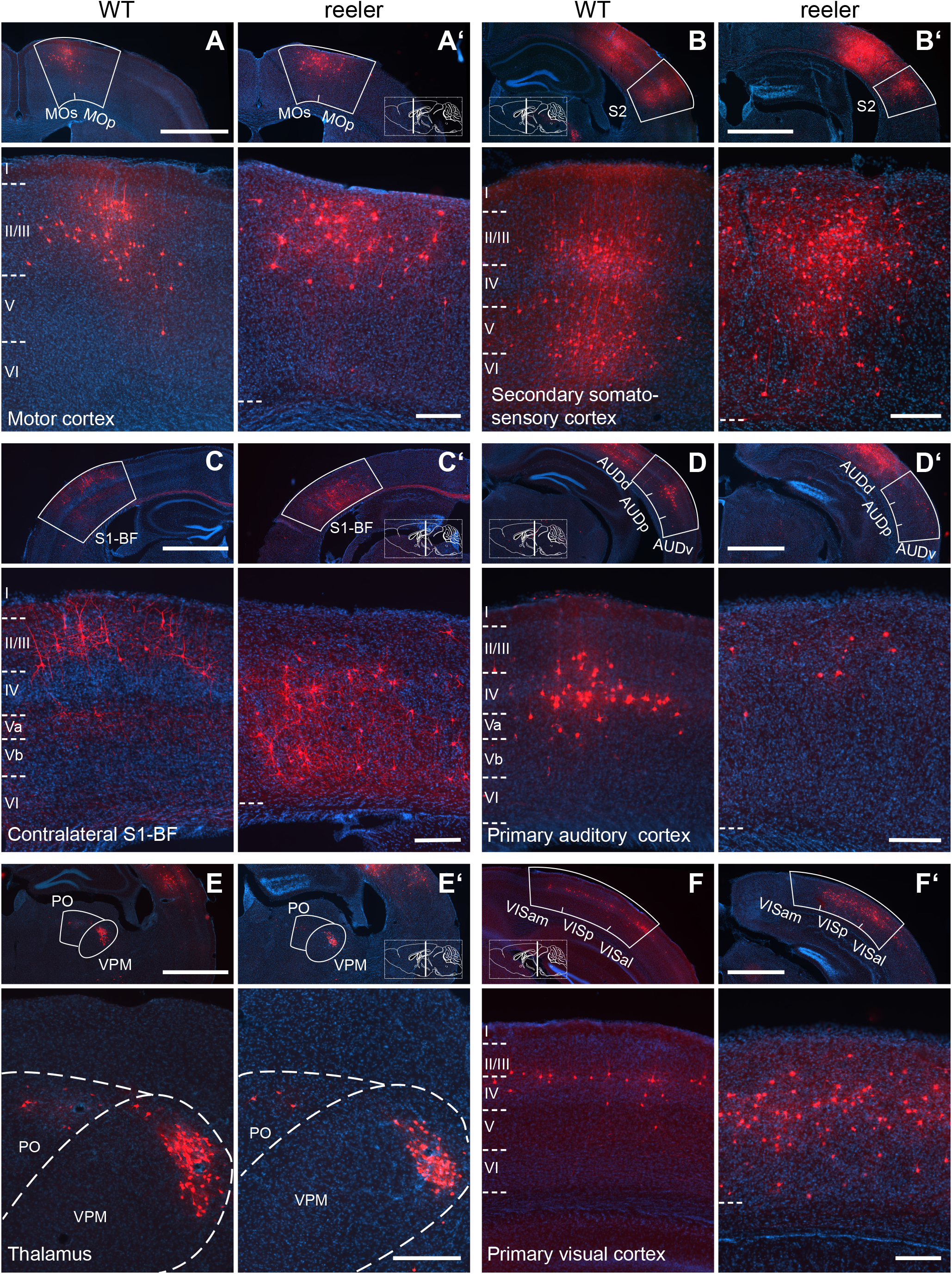
Long-range input to VIP cells in barrel cortex of WT and reeler mice. (A-F, A’-F’) Sections along the rostro-caudal extent of WT and reeler mice showing consistently labeled areas with presynaptic partners of VIP cells in barrel cortex. In the top overview panels, the white contour delineates the borders of the respective area. Higher magnification close-ups are shown below. Section planes are indicated on the schematic sagittal brain section. (scale bar overview: 1000 μm; scale bar close-up: 200 μm; Abbr.: AUDd/AUDp/AUDv, dorsal/primary/ventral auditory area; MOp/MOs, primary/secondary motor cortex; PO, posterior complex of the thalamus; S1-BF, primary somatosensory cortex, barrel field; S2, secondary somatosensory cortex; VISal/VISam/VISp, anterolateral/anteromedial/primary visual area; VPM, ventral posteromedial nucleus of the thalamus).

### VIP cells in reeler displayed an imbalance between ipsi- and contralateral input

When analyzing quantitative differences in the inputs between genotypes, we first focused on more global categories of input (Figure 4, Table S1). We first summed up all long-range inputs and then separated inputs from ipsilateral and contralateral cortical areas, from the thalamus and from all subcortical areas. We normalized the inputs by dividing the cell count by the number of starter cells to calculate the input magnitude, which can be seen as a proxy for the strength of an input. VIP neurons in reeler received overall fewer inputs per cell. This reduction was exclusively due to fewer inputs from the ipsilateral cortex in reeler, while they received more contralateral inputs per cell. There was no difference for thalamic or total subcortical input.

**Figure 4:**
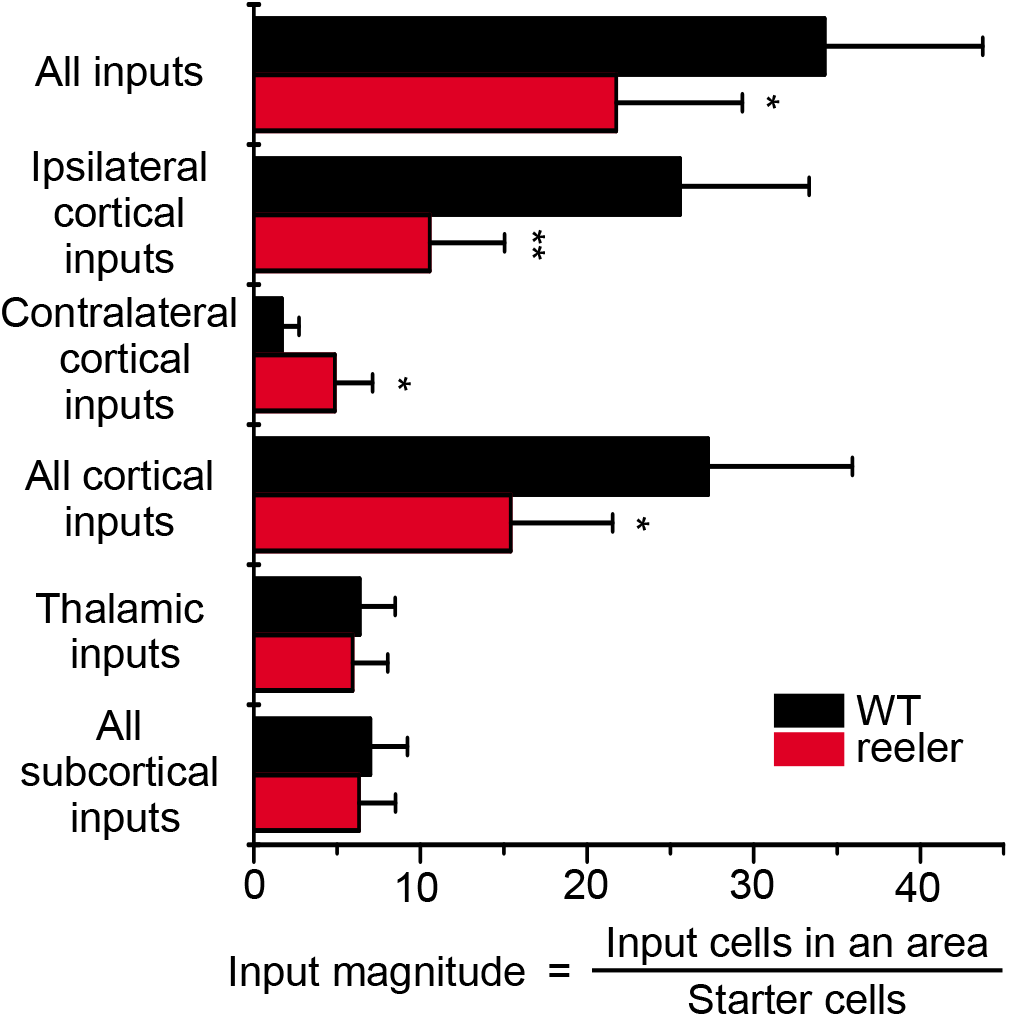
Input magnitude from global cortical and subcortical areas highlight a cortical phenotype. Histograms representing the input magnitude from summed-up cell counts of brain areas. Reeler mice received overall less input per cell, which was due to less input from the ipsilateral cortex. Input from the contralateral hemisphere was increased in reeler whereas subcortical input remained unaffected. (n = 7 per group; mean ± SD; *p < 0.05, **p < 0.01).

The input magnitude might be influenced by the number of starter cells. One presynaptic neuron could synapse on multiple starter cells. This divergent connectivity prompts that with a rising number of starter cells, the count of additionally labeled postsynaptic cells would decrease. Hence, the higher the number of starter cells, the lower the ratio of starter cells to input cells (input magnitude). The smaller overall input magnitude in reeler might result from this group having on average more starter cells. To investigate this relationship we plotted the number of starter cells against the input magnitude for the two genotypes (Figure S2C). For both genotypes the slope was close to zero (WT: Input magnitude = 0.002*starter cells+33.9, R^2^= 0.2; reeler: Input magnitude = −0.002*starter cells+22.2, R^2^= −0.2). In consequence, there was no relationship between the number of starter cells and the input magnitude, and hence the higher number of starter cells in reeler cannot explain the lower input magnitude.

In sum, VIP neurons in reeler mice are embedded in a different long-range cortical circuitry that contains fewer inputs from the ipsilateral and more inputs from the contralateral cortical hemisphere.

### Auditory and motor cortex showed the strongest reduction in ipsilateral input

Because each VIP neuron in reeler received on average less input, the subsequent analyses for differences among individual areas using the input magnitude would be inherently biased. Therefore, we employed another means of normalization: input fraction. It is the number of inputs in an area divided by the number of total inputs in the brain. It reflects how different inputs are balanced. We selected 41 consistently labeled areas that constituted the majority of inputs (97.6% in WT; 98.8% in reeler). We calculated the input fraction for each area and made pairwise comparisons between genotypes (Figure 5, Table S2). For motor, somatosensory body region, auditory and visual cortex we also summed up counts in the individual sub-areas to calculate a total input fraction. VIP neurons in reeler mice received a notably lower fraction from almost all ipsilateral cortical areas. The strongest reduction was seen for primary auditory cortex. The input fraction for motor cortex was also significantly reduced. The contralateral barrel cortex as well as the contralateral secondary somatosensory cortex contributed a significantly higher fraction of inputs in reeler. Subcortical input fraction was fairly similar and only the ventral posterolateral nucleus of the thalamus constituted a higher fraction of inputs in reeler. This analysis of the fraction of inputs validates that the proportion of inputs from the ipsilateral hemisphere is reduced and from the contralateral homotopic area increased.

**Figure 5:**
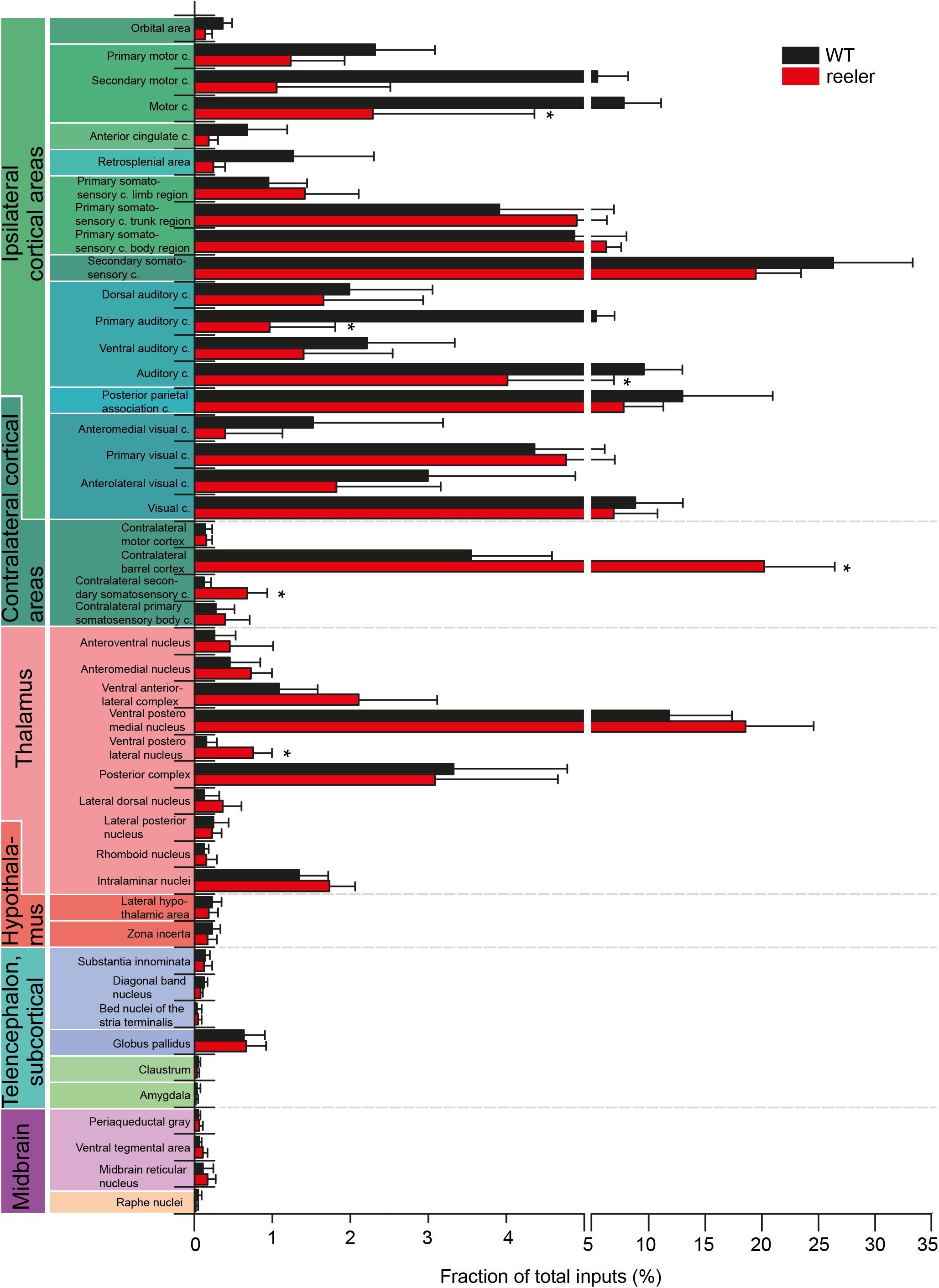
Comparative analysis of the fraction of inputs from individual areas. Mean proportion of RV-labeled cells in 41 individual areas normalized against the total number of inputs in the whole brain for the two genotypes. For motor cortex, primary somatosensory cortex body region, auditory cortex and visual cortex, the summated proportions of the individual subareas are shown as well. Pairwise comparisons were carried out to assess differences in input fraction for individual areas. For individual values see Table 1 (n= 7 per group; mean ± SD; *p < 0.05; Abbr.: c, cortex).

**Table 1:** Mean and standard deviation (SD) for input magnitude from global areas. P-Values were calculated with pairwise comparisons and adjusted for multiple comparisons. Related to Figure 5.

| Area | WT |  | Reeler |  | p-value | adj.p-value |
| --- | --- | --- | --- | --- | --- | --- |
|  | mean | SD | mean | SD |  |  |
| All inputs | 34.26 | 9.49 | 21.74 | 7.58 | 0.0183 | 0.0183* |
| Ipsilateral cortical inputs | 25.59 | 7.74 | 10.56 | 4.49 | 0.0008 | 0.0032** |
| Contralateral cortical inputs | 1.68 | 1.03 | 4.86 | 2.26 | 0.0175 | 0.0355* |
| All cortical inputs | 27.27 | 8.64 | 15.42 | 6.11 | 0.0118 | 0.0355* |
| Thalamic inputs | 6.36 | 2.13 | 5.93 | 2.10 | 0.7124 | 1.0000 |
| All subcortical inputs | 6.99 | 2.24 | 6.32 | 2.18 | 0.5819 | 0.5819 |

**Table 2:**
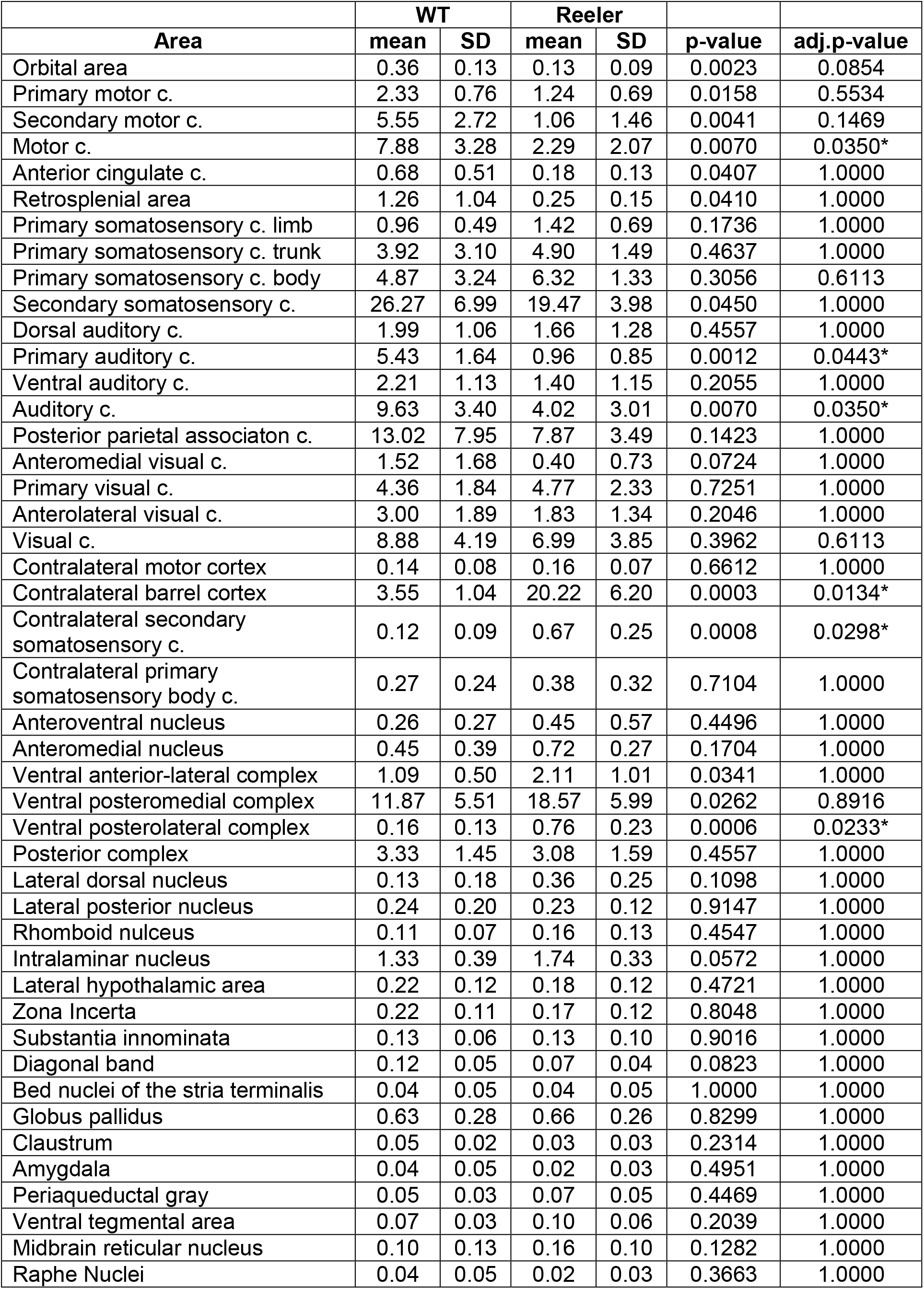
Mean and standard deviation (SD) for input fraction from individual areas. P-Values were calculated with pairwise comparisons and adjusted for multiple comparisons. Related to Figure 6.

### Reeler mice had a larger corpus callosum

The fact that VIP cells in reeler received more input from the contralateral somatosensory cortex prompted the question if there are more callosal projection neurons (CPNs) in reeler. We injected the retrograde tracer AAV-retro-EGFP (Tervo et al., 2016) into the right barrel cortex of WT and reeler mice and counted the CPNs in the other hemisphere (Figure S4A, A’). We normalized the count of CPNs by the number of labeled cells in a given volume at the injection site. In reeler there was about a 3-fold increase in the average number of CPNs (Figure S4C). This increase was the same as for contralateral inputs to VIP cells. Moreover, the distribution of CPNs was very similar to the contralateral inputs’ to VIP cells, being more biased towards the lower part of the cortex (Figure S4D, Figure 6F).

**Figure 6:**
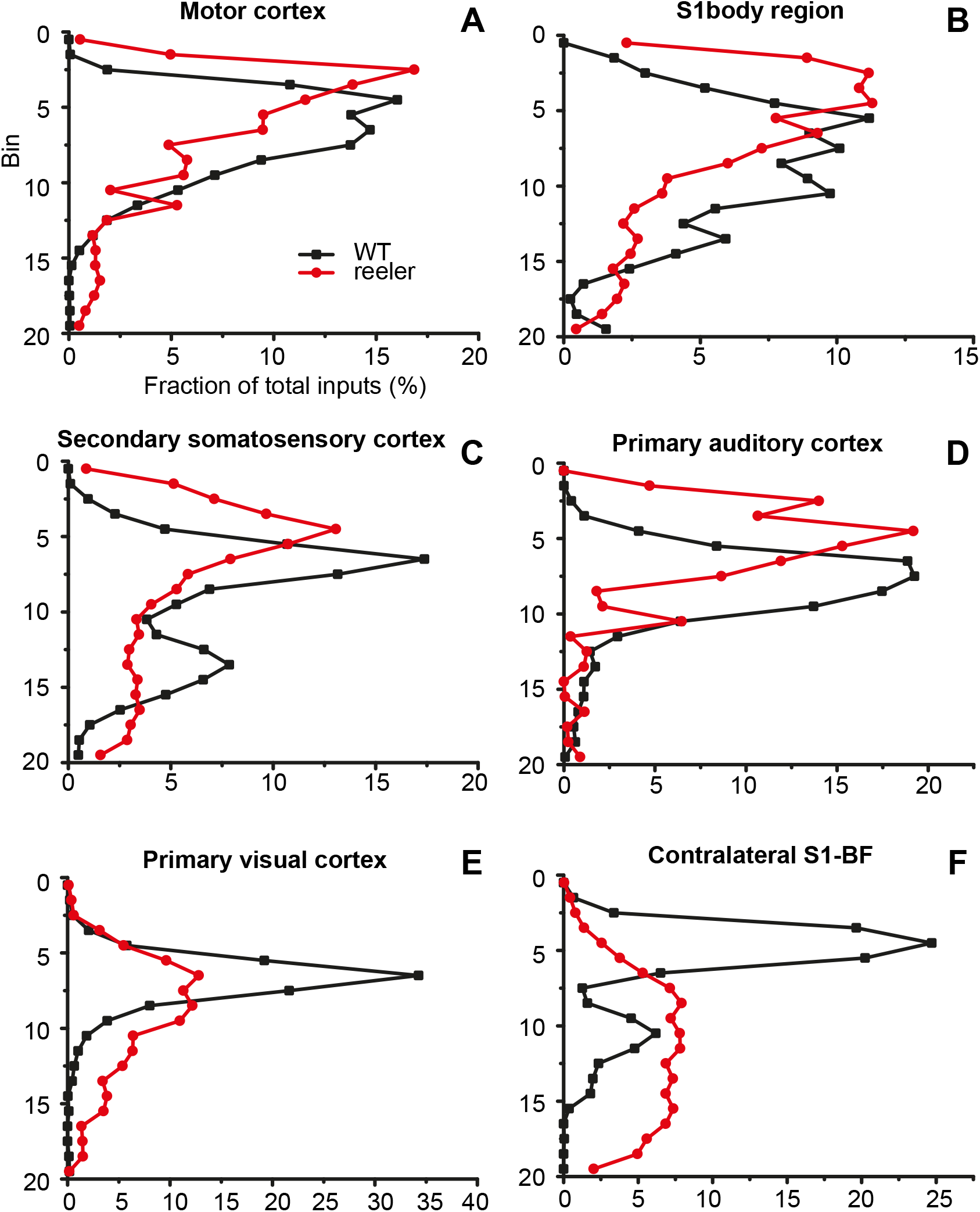
Distribution of projection neurons in cortical input areas. (A-F) The thickness of the cortex was divided into 20 equal-sized bins. We plotted the fraction of inputs in a bin normalized against the total inputs of VIP cells from this area. In ipsilateral areas, the distribution of projection neurons had one peak that was usually shifted more superficially in reeler, except for visual cortex where the peak was just broader. The distribution of contralateral projection neurons from the barrel cortex was very different, with neurons in WT mostly in the upper third, while in reeler predominantly in the lower two thirds of the cortex (Abbr.: S1, primary somatosensory cortex; BF, barrel field).

If there are more CPNs in reeler, the corpus callosum should also contain more fibers and hence be larger. To provide additional evidence for an increased callosal connectivity, we performed magnetic resonance imaging (MRI) to measure the dimensions of the corpus callosum. We performed a quantitative morphometric analysis on three WT-reeler littermate pairs. We segmented the corpus callosum based on structural information of magnetization transfer maps (Figure S5A, A’) and directional information of fractional anisotropy maps (Figure S5C, C’). We measured the area of the corpus callosum on the mid-sagittal section, which was larger in reeler in every littermate pair (Figure S5D). This effect was not generated by differences in cortical volume as the volume of the isocortex was fairly well matched between littermates, although in one pair the volume was slightly larger in the reeler mouse (Figure S5E). Furthermore, we noticed on coronal sections that the corpus callosum was shaped differently in reeler. It did not have a typical U-shape at the medial area but was flattened out (Figure S5B, B’). These experiments indicate that reeler mice have a larger corpus callosum because of a surplus of CPNs sending an axon to the other hemisphere.

### Ipsilateral projection neurons were distributed similarly but CPNs differently

In reeler mice, the whole cortex shows alterations because of reelin deficiency. Therefore, the presynaptic projection neurons labeled in our study should show a markedly different distribution than in WT. To investigate the pattern of dispersion, we divided the cortex into 20 equal-sized bins and counted the proportion of presynaptic cells in each bin for ipsilateral motor, primary visual, primary auditory, somatosensory body (trunk+limbs) and secondary somatosensory cortex as well as for contralateral barrel cortex (Figure 6). In most ipsilateral areas, projection neurons in reeler mice had a very similar distribution across the cortical depth compared to WT but slightly shifted towards the pial surface (Figure 6A-D). Only in visual cortex the distribution looked notably different. In WT, most projection neurons were located around the LIII-IV border. This peak was smoothed out in reeler, indicating that the projection neurons were rather dispersed than inverted in their arrangement (Figure 6E). The pattern of projection neurons in the contralateral barrel cortex, however, was completely different (Figure 6F). In WT, projection neurons were predominantly located in the upper third of the cortex, corresponding to LII/III. In reeler, projection neurons were predominantly located in the lower two-thirds of the cortex.

## DISCUSSION

The main aim of this study was to investigate if the distribution of cortical cells across layers is linked to their capacity to integrate long-range inputs. We investigated the long-range inputs to VIP neurons because they have a very typical laminar bias towards the upper layers, which is lost in reeler mice. Using retrograde RV-tracing we mapped the brain-wide inputs to VIP neurons in barrel cortex of WT and reeler mice. There was no difference in the innervation by subcortical inputs. Cortical inputs were from the same sources, however, in reeler ipsilateral cortical inputs were reduced and contralateral cortical inputs were much more numerous, revealing a different cortical circuit architecture in the absence of layers.

### Input to VIP cells originated from the same sources in WT and reeler

We found that VIP cells in the barrel cortex of WT and reeler mice were innervated by the same sources in both genotypes. This is not surprising because the existence of the same types of projection neurons has been confirmed in reeler, although in cortex they are in ectopic positions (Caviness, 1976; Diodato et al., 2016; Imai et al., 2012; Steindler and Colwell, 1976; Wagener et al., 2016; Yoshihara et al., 2010). Numerous tracing studies of cortical neurons in WT mice, including excitatory (DeNardo et al., 2015) and inhibitory (Wall et al., 2016) neurons in barrel cortex, demonstrated that all neuron classes in a cortical area receive input from the same sources. Therefore, the feature, which is considered to distinguish cell types and relate to potential functional implications, is the input’s proportion a given neuron type receives. In consequence, we expected to discover putative differences in the afferent connectivity to VIP neurons between genotypes specifically on a quantitate level, which has not been assessed for long-range projections in reeler yet.

### Subcortical fibers target VIP cells independent of laminar position

When we looked at the proportion of inputs to VIP cells, we found profound differences between WT and reeler. VIP cells in reeler received less input per cell than in WT, which was exclusively due to a reduction in cortical inputs, while the subcortical inputs remained the same.

The thalamic input comprised the main source of subcortical input. The cells in the thalamus are at most only subtly affected by the reelin mutation (Lambert de Rouvroit and Goffinet, 1998; Wagener et al., 2010). Previous studies have shown that thalamic fibers from the VPM reach their malpositioned postsynaptic targets in the reeler cortex although they take an unusual trajectory, reaching LI before bouncing back to plunge down on the target cells (Harsan et al., 2013; Molnár et al., 1998; Wagener et al., 2016). Together with our results, these studies support the idea that fibers from properly developed subcortical structures find their ectopic targets in the mislaminated reeler cortex in approximately the same numbers. In consequence, the laminar distribution of VIP cells seems not to be linked to their capacity to integrate subcortical long-range inputs.

### Reeler mice have increased callosal connectivity

In the reeler cortex, both the VIP cells as well as their cortical presynaptic partners are malpositioned. In this case, we found the circuit organization to be different. The average number of ipsilateral long-range projection neurons converging on a VIP cell was significantly lower than in WT. Especially the proportion of inputs from motor and auditory cortex to VIP cells in barrel cortex was strongly reduced.

In parallel to the reduction of ipsilateral cortical input, there was a massive increase of afferents from the contralateral hemisphere. We showed that in reeler there are more CPNs connecting the hemispheres of barrel cortex and as a result the corpus callosum is larger. Although previous studies have investigated the macroanatomy and trajectory of fiber bundles in the reeler mouse (Badea et al., 2007; Harsan et al., 2013), the enlargement of the corpus callosum has remained unnoticed. Similarly, earlier tracing studies have just confirmed the existence of callosal connections between homotopic areas but did not assess the numbers of CPNs (Caviness and Yorke, 1976; Imai et al., 2012; Steindler and Colwell, 1976). At least, our observation that the distribution of CPNs is biased towards deeper parts of the cortex is not without precedent (Steindler and Colwell, 1976).

### Different mechanisms might drive the formation of ipsi- and contralateral projections

Because ipsi- and contralateral long-range inputs showed such an opposing change in innervation intensity, we speculate that these two types of connections mature at different time points and based on different mechanisms. A sizeable fraction of ipsilateral projection neurons in reeler either fail to grow an axon or grow an axon but fail to find their destined postsynaptic partner. There is subtle evidence that ipsilateral projection neurons do not form excess projections that are pruned away but have a directed outgrowth and their connections are stably established by postnatal day (P)7 (Klingler et al., 2018). On the contrary, during the development of callosal projections a transiently higher number of callosal axons is reduced in an activity-dependent process of axonal elimination (rev. in (Innocenti and Price, 2005)). In the somatosensory system, CPNs invade the contralateral cortex by P5, reach their maximum density by P10 and then reduce again to reach a stable level of innervation by P20 (Fenlon et al., 2017; De Leon Reyes et al., 2019). Because the establishment of permanent callosal synapses is only finished in the third postnatal week, they might mature based on the network activity generated by already existing ipsilateral connections (Petreanu et al., 2007; Suárez et al., 2014). Perhaps in reeler, the pruning of callosal axons happens to a lesser degree than in WT to maintain a stable network in which the inputs of multiple afferent systems are balanced (Caviness and Rakic, 1978).

Another possibility would be that the absence of reelin impacts signaling pathways for axonal outgrowth. For example, Sema3a is a signaling molecule involved in axonal guidance that acts as a repulsive cue for axons (Polleux et al., 1998). Knock-out of the sema3a receptor causes an increase in contralateral axon projections (Wu et al., 2014). Sema3a and reelin pathways could intersect at the Fyn kinase, which appears in the intracellular signaling cascade of both factors (Lee and D’Arcangelo, 2016; Sasaki et al., 2002). Indeed, Fyn deficiency causes a higher maintenance rate of growth cones (Sasaki et al., 2002) and knock-out mice show a lamination defect (Kuo et al., 2005). Perhaps the absence of reelin collapses the synergy of these two pathways and leaves more callosal axons intact.

### An alternative circuit to retain cognitive abilities

Despite the absence of layers, reeler mice display no severe decline in cognitive abilities, have largely preserved sensory function, and only have a slight impairment of spatial memory and executive function (Imai et al., 2017; Pielecka-Fortuna et al., 2015; Salinger et al., 2003; Wagener et al., 2010). So far this preservation of cognitive abilities has been attributed to the fact that in reeler neurons seem not only to retain their physiological properties but also their connectivity, implying the formation of the same circuits despite the malposition of its parts (Guy and Staiger, 2017). However, our map of brain-wide inputs to VIP neurons reveals that reeler mice have a different proportion of ipsi- and contralateral input. We argue that this change in connectivity might emerge as a plastic adaption in pursuit of retaining global cognitive abilities like sensory perception but perhaps fails to realize more specialized and challenging domains of cognition. Because our connectivity map contains information on the number of connections, it allows to make very specific assumptions about behavioral impacts. For example, the auditory cortex fuels information about sound to the somatosensory cortex to be integrated into tactile processing (Lemus et al., 2010; Maruyama and Komai, 2018). This input pathway comprises much fewer neurons in reeler. Behavioral experiments compelling reeler mice to integration of sound into tactile perception might be a starting point to study the functional significance of a diminished input. Similarly, the increased callosal connectivity might hint that reeler mice rely more on bilateral information during their sensory exploration. This could be tested perhaps in a variation of a corridor tracking task adapted for reeler to exclude influence of motor deficits (Sofroniew et al., 2015).

In conclusion, we incline to the viewpoint that layers are important for the formation of the mouse cortical network as we know it, maybe providing the guidance for pre- and postsynaptic partners to meet across long distances. However, there are plastic mechanisms independent of neuronal position allowing alternative circuits to form that enable basic cognition. Looking across species, intelligence presents itself in various connectivity schemes. The analogue to the mammalian cerebral cortex in birds is the pallium, which is organized in nuclei (Butler and Cotterill, 2006). The analogue in cephalopods is the vertical lobe, which is organized in serial matrices (Shigeno et al., 2018). Both taxa evolved species with baffling cognitive abilities rivaling mammalian intelligence (Roth, 2015). There is even strong evidence that in the bird pallium neurons express the same molecular markers and assemble the same microcircuits found in mammals despite their different arrangement (Calabrese and Woolley, 2015; Dugas-Ford et al., 2012). Looking at different wiring approaches will benefit our understanding of the essential features neuronal circuits require to give rise to cognitive functions.

## ACKNOWLEDGEMENTS

This work was supported by the Deutsche Forschungsgemeinschaft via CRC 889 (Cellular mechanisms of sensory processing; TP C07 to J.F.S.) and STA 431/11-2.

We thank Patricia Sprysch and Sandra Heinzl for excellent technical assistance. We are grateful to Karl-Klaus Conzelmann for kindly donating RV-SADΔG-mCherry (EnvA). We thank Michael Lingelbach and Edward Callaway for kindly donating pAAV-DIO-TVA^66T^-EGFP-oG.

## AUTHOR CONTRIBUTIONS

J.F.S., J.G. and G.H. conceived the study. G.H. conducted and analysed all tracing experiments. J.G. and M.W. gave experimental advice. P.T. performed microscopic imaging. A.R. analysed the VIPCre/reeler/tdTomato mouse line. N.S, R.D. and S.B. performed the MRI experiments. G.H., N.S and J.F.S. wrote the original draft.

---

**Figure S1:**
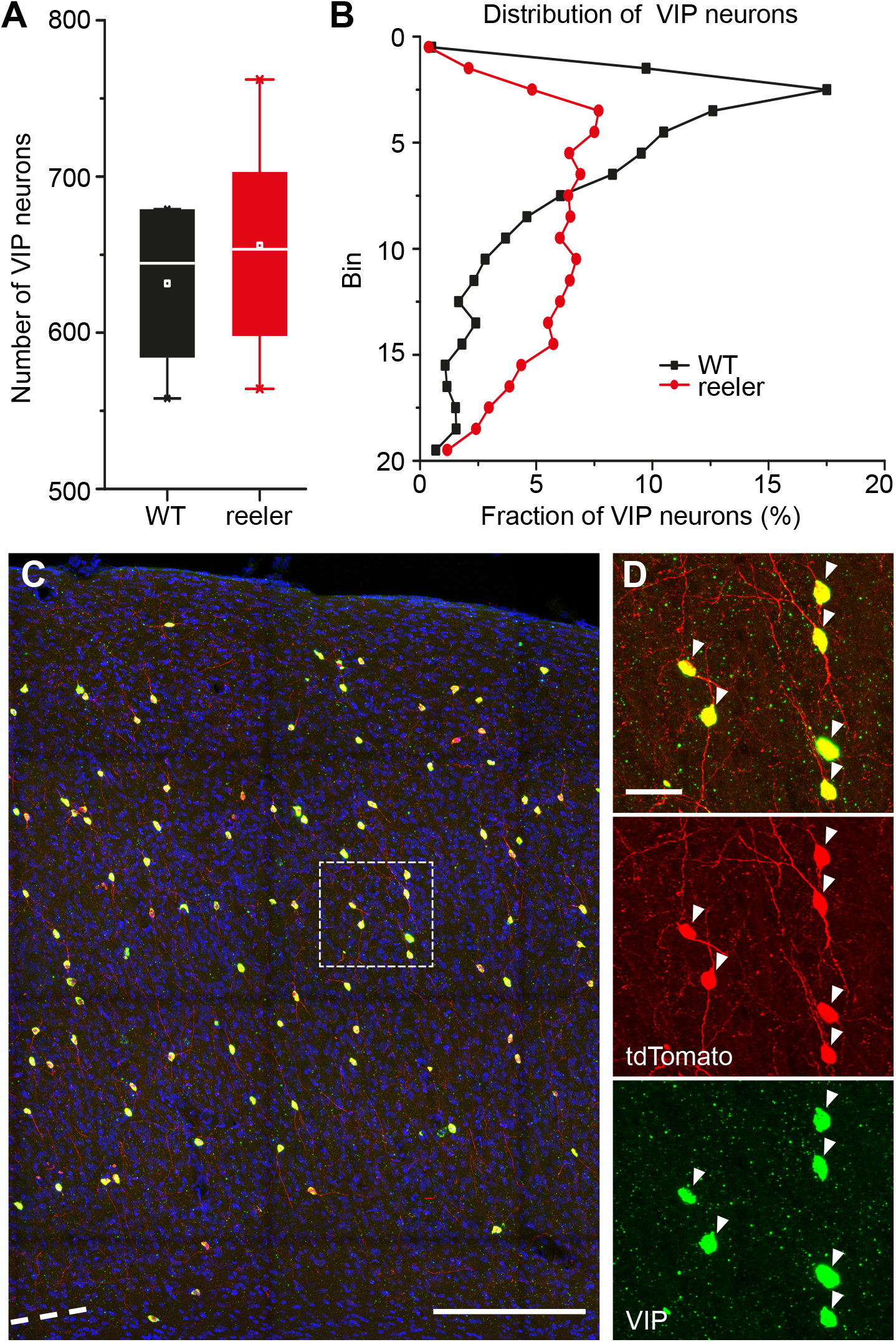
VIP neurons in the reeler mouse retain numbers and VIP expression but lose laminar bias. Related to Figure 1. (A) Number of VIP neurons counted in a volume of cortex that spanned from pia to white matter and was 100 μm wide and 240 μm thick (n=5 WT mice; n=4 reeler mice; box plot: white line = median; white dot = mean). Counts were almost the same in WT and reeler. (B) Distribution of VIP neurons across the cortical depth. In WT they showed a prominent peak in the upper layers. In reeler they were fairly uniformly distributed with few neurons close to pia and white matter. (C) Section of barrel cortex of a VIP-Cre/reeler/tdTomato mouse stained against VIP-RNA (green) with fluorescent in-situ hybridization. Virtually all tdTomato cells overlapped with VIP signal (scale bar: 200 μm). (D) Insert of C with individual channels to reveal co-localization of tdTomato and VIP signal (scale bar: 20 μm).

**Figure S2:**
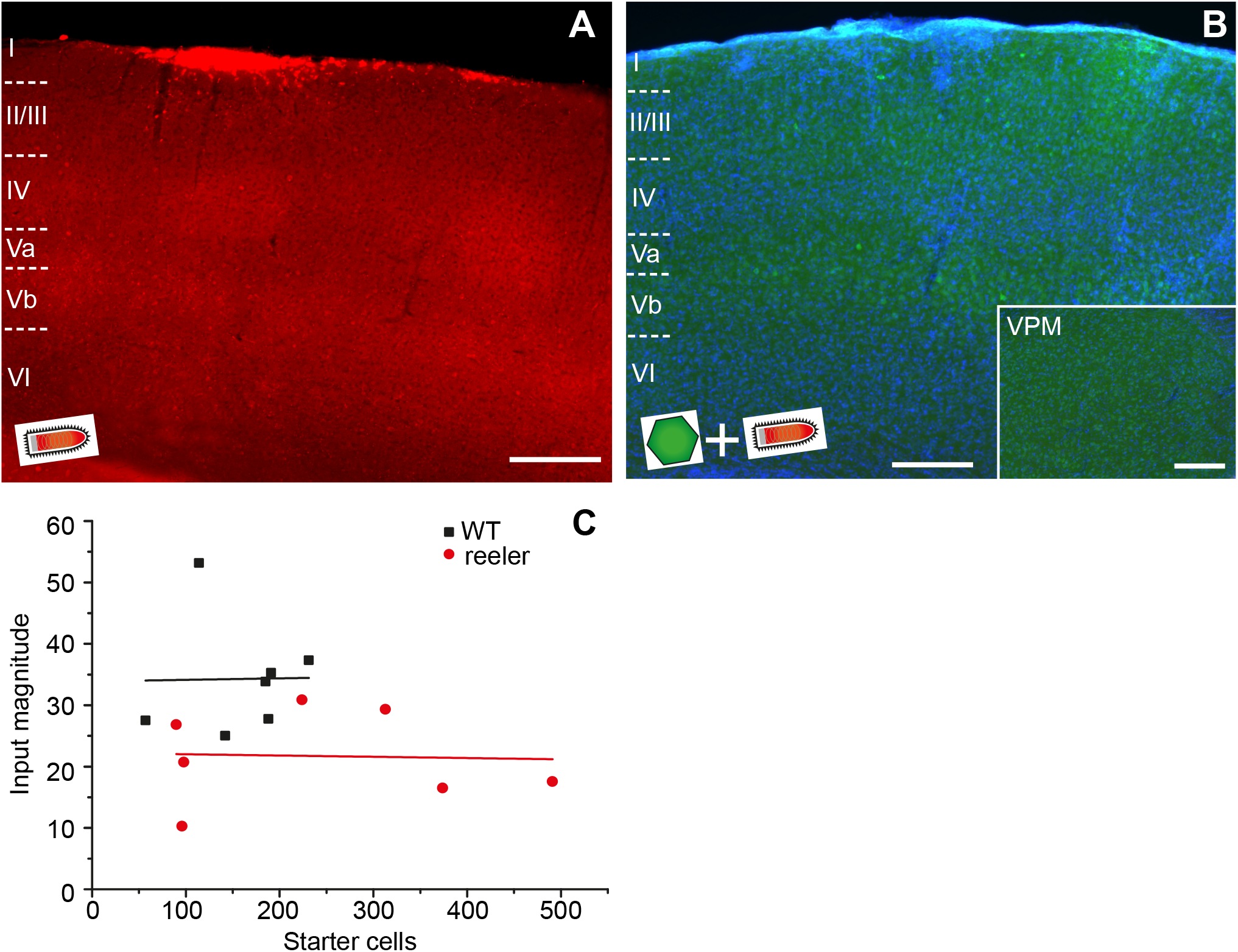
Validation of constructs for RV-tracing in BL6 animals. Related to Figure 2. (A) Coronal section through the barrel cortex of a BL6 mouse (wild type) after injection of RV-mCherry, without prior injection of helper AAV. No transduced cells were detected. Uptake of RV into cells strictly depended on the presence of TVA (scale bar: 200 μm). (B) AAV8-DIO-TVA66T-EGFP-oG and RV-mCherry were injected with the same titer as in experimental conditions but in a BL6 (wild type) animal. We did not observe any RV labeling at the injection site nor in the ventral posteromedial nucleus of the thalamus (insert), a structure with reliable input to the barrel cortex. Therefore, this AAV does not show leak expression in the absence of Cre that would confound the tracing experiments (scale bar: 200 μm). (C) Input magnitude was plotted against starter cell count for WT and reeler. There was no correlation between the two variables. Therefore, the difference in input magnitude between genotypes cannot be explained by a difference in starter cell numbers.

**Figure S3:**
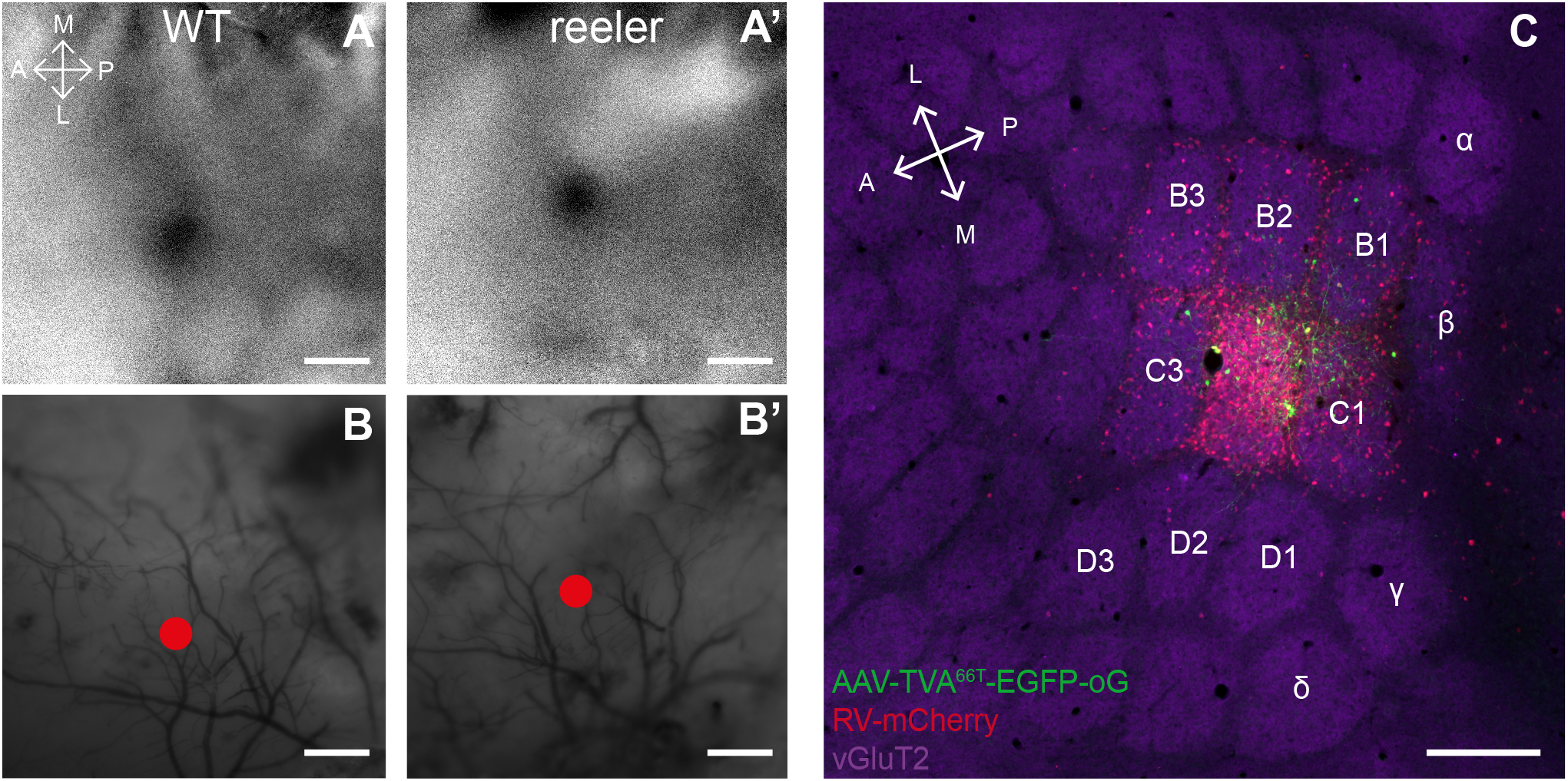
Mapping the whisker C2-related cortical module in barrel cortex to coherently target the tracer injections. Related to Figure 2. (A, A’) Red light was shone on the exposed cortical surface and its reflectance was measured with a CCD-camera, while the contralateral C2 whisker was stimulated. Repetitive whisker stimulation over 30 trials led to a localized change in blood flow, which induced a change in light reflectance visible as a dark spot. WT and reeler mice had very similar signals in terms of dynamics and size (scale bar: 200 μm). (B, B’) Surface vasculature was overlaid with image in A and the location of the strongest change in reflectance was marked. The blood vessels were used as landmarks to guide the injection pipette to the dot (scale bar: 200 μm). (C) Tangential section of barrel cortex after targeted injection of AAV-TVA66T-EGFP-oG and RV-mCherry into the C2 column. Staining thalamic terminals with vesicular glutamate transporter 2 (vG-LuT2) allowed to visualize the barrels. The density of input cells was highest in C2 indicating that the majority of VIP starter cells was located within this barrel-related column (scale bar: 200 μm).

**Figure S4:**
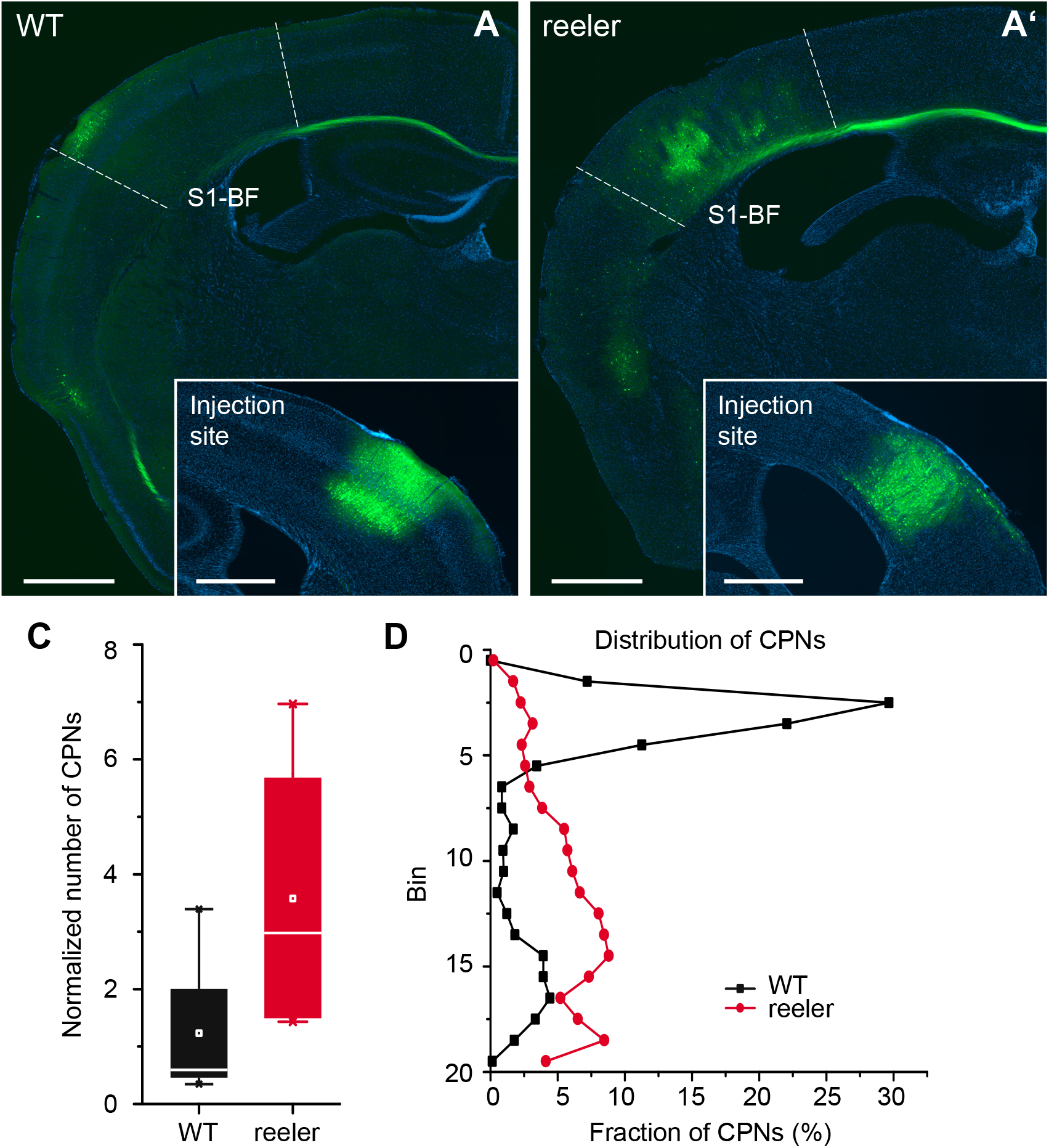
Retrograde tracing of callosal projection neurons (CPNs). Related to Figure 4. (A, A’) Coronal sections through the barrel cortex contralateral to the injection site of AAV-retro-EGFP. CPNs are more numerous and more dispersed in reeler. The injection site is shown in the insert (scale bar: overview: 1000 μm; insert 500 μm). (B) Number of CPNs normalized by number of cells at the injection site in a volume from pia to white matter with a 200 μm diameter around the injection site. Reeler mice had about 3 times more CPNs than WT. (D) Distribution of CPNs across the cortical depth. The cortex was divided into 20 bins of equal size and the relative fraction of CPNs in each bin was plotted. CPNs in WT were predominantly in the upper part corresponding to LII/III. In reeler they were more in the deeper parts of the cortex. The distribution of all CPNs wes similar to the distribution of CPNs targeting VIP cells in both genotypes.

**Figure S5:**
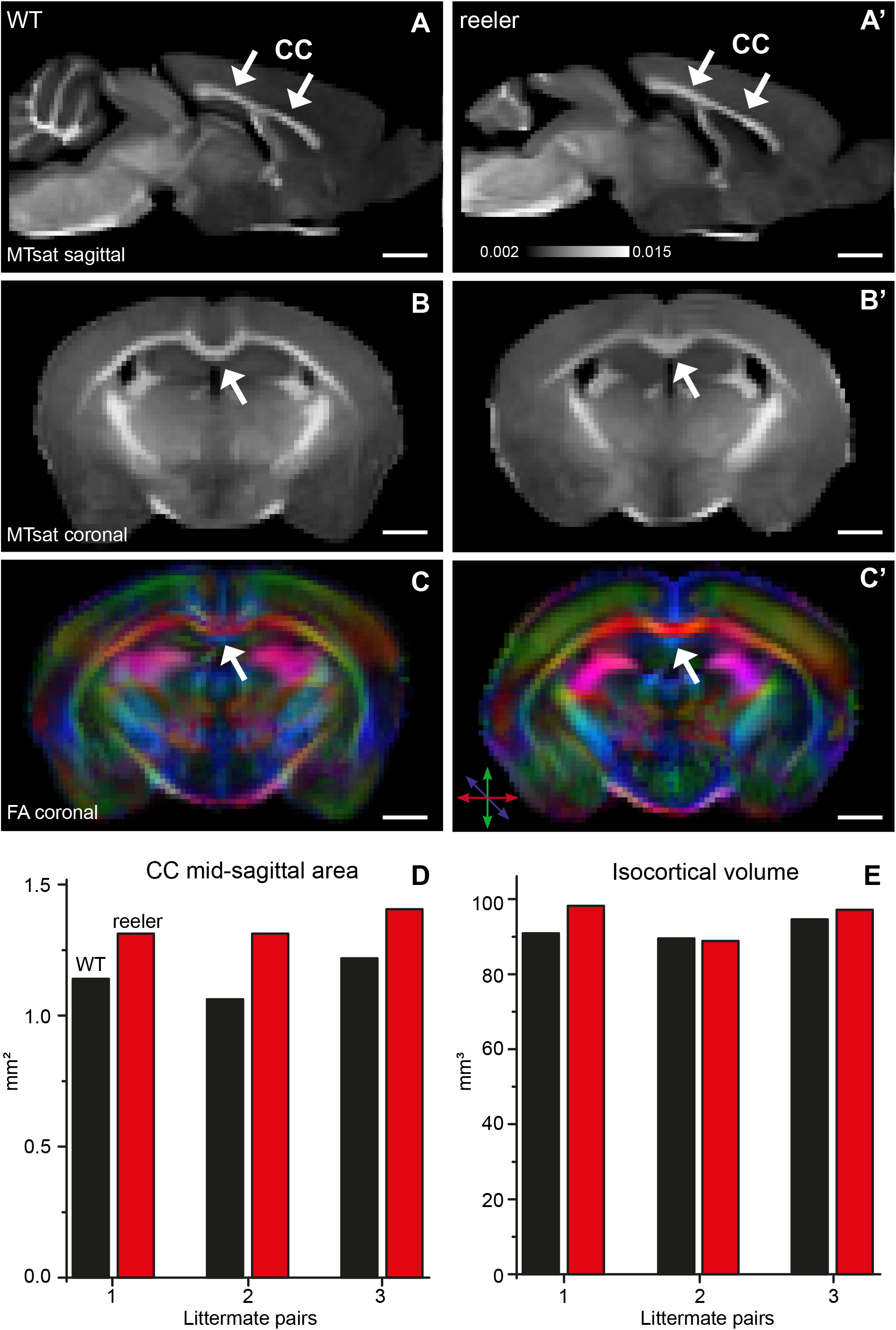
MRI of WT and reeler mice to assess dimensions of corpus callosum. Related to Figure 4. (A, A’) Sagittal maps of magnetization transfer saturation (MTsat) of a WT and reeler mouse brain acquired ex-vivo. These maps were used to determine the dimensions of the corpus callosum (CC). (B, B’) Coronal MTsat map of WT versus reeler mouse. At the midline the CC showed a different geometry between genotypes. In reeler mice the characteristic curvature of WT mice was absent so that the top of the CC appeared flatter. (C, C’) Fractional anisotropy (FA) maps show the color-coded directionality of fibers. Because fibers of the CC run in the medio-lateral direction they could be distinguished from whiter matter bundles more dorsally or more ventrally that run in the rostro-caudal direction. (D) Mid-sagittal area of the CC of individual WT and reeler mice compared between littermate pairs. In each pair, the reeler mouse had a larger area, probably indicating that it is comprised of a higher number of callosal fibers. (E) Total isocortical volume of individual WT and reeler mice compared between littermate pairs. The volume was almost the same except for one pair in which the reeler mouse showed a slightly higher volume.

